# Comparative Analysis of Denitrification and Other Nitrogen Cycle Genes in Diverse Environmental Microbiomes

**DOI:** 10.1101/2025.08.10.669474

**Authors:** Andrew M. Pilat, Rachael E. Tomasko, Audrey Gibson, Daniel Barich, Zachary Slimak, M. Siobhan Fennessy, Joan L. Slonczewski

**Author notes:** Corresponding author: Joan L. Slonczewski, Department of Biology, Kenyon College, Gambier, OH.

## Abstract

The nitrogen cycle includes the microbial transformation of reactive nitrogenous molecules within a given ecosystem. Shotgun metagenomics, the unbiased sequencing of a microbial community, can reveal the unique gene profile and community composition of diverse microbiomes. We used NCycDB, an assembly-free, alignment-based pipeline, to determine and compare the potential nitrogen cycle processes in seven distinct microbiomes: wetland soils, agricultural soils, forest soils, compost, river water, and pond water from Knox County, Ohio, and lake water from the McMurdo Dry Valley Region in Antarctica. Soil metagenomes showed high levels of bacterial denitrifier genes, whereas freshwater metagenomes showed fewer denitrifiers. In wetland and agricultural soil cores, the relative abundance of *norB*, the marker gene for nitric oxide reductase, was a predictor of ambient N_2_O flux as measured by gas chromatography. Denitrifiers were predicted using the Kraken2/Bracken pipeline. In Ohio wetland soils, the relative abundance of *Bradyrhizobium* species were strongly associated with denitrification genes. We also explored how freshwater environmental factors select for or against nitrogen cycle genes. Denitrifier genes were positively correlated with phosphate concentration. Denitrifiers and denitrification genes were detected under microaerobic conditions, demonstrating that denitrification activity diminishes proportionally to oxygen concentration. An improved understanding of the denitrifier community in diverse microbiomes is necessary to mitigate excess N_2_O emissions from anthropogenic activities such as wetland drainage and crop production.

**IMPORTANCE:** Terrestrial nitrogen limitations have historically reduced crop output. To fertilize crops, humans have developed synthetic N fixation methods, such as the Haber-Bosch process, which now account for half of global nitrogen uptake in biomass. The resulting nitrogen surplus in soils leads to eutrophication, harmful algal blooms in lakes and other aquatic ecosystems, and increased N_2_O emissions from denitrification via anaerobic respiration by soil bacteria. Considering that the nitrogen cycle is controlled by microbial communities, we need a better understanding of how nitrogen-cycling microbes control N runoff and N_2_O flux in each unique microbiome.

## INTRODUCTION

Despite being the most abundant element in the atmosphere, nitrogen (N) is commonly the limiting reagent for any given terrestrial ecosystem (1). The invention of the Haber-Bosch process in 1909 revolutionized fertilizer production but also led to huge imbalance in nitrogen cycle and accelerated our dependence on petroleum-based energy (2–4). A staggering 83% of fertilizer nitrogen is oxidized by nitrifying bacteria, forming nitrites and nitrates (**Figure 1**) (3). Nitrate is highly water soluble and flows into waterways as agricultural runoff, promoting harmful algal blooms (HAB) (5,6). Over 20 percent of US shallow water tables in agricultural regions exceed federal nitrate concentration standards, which could increase the risk of cancer, diabetes, and miscarriage (7,8).

**Figure 1.**
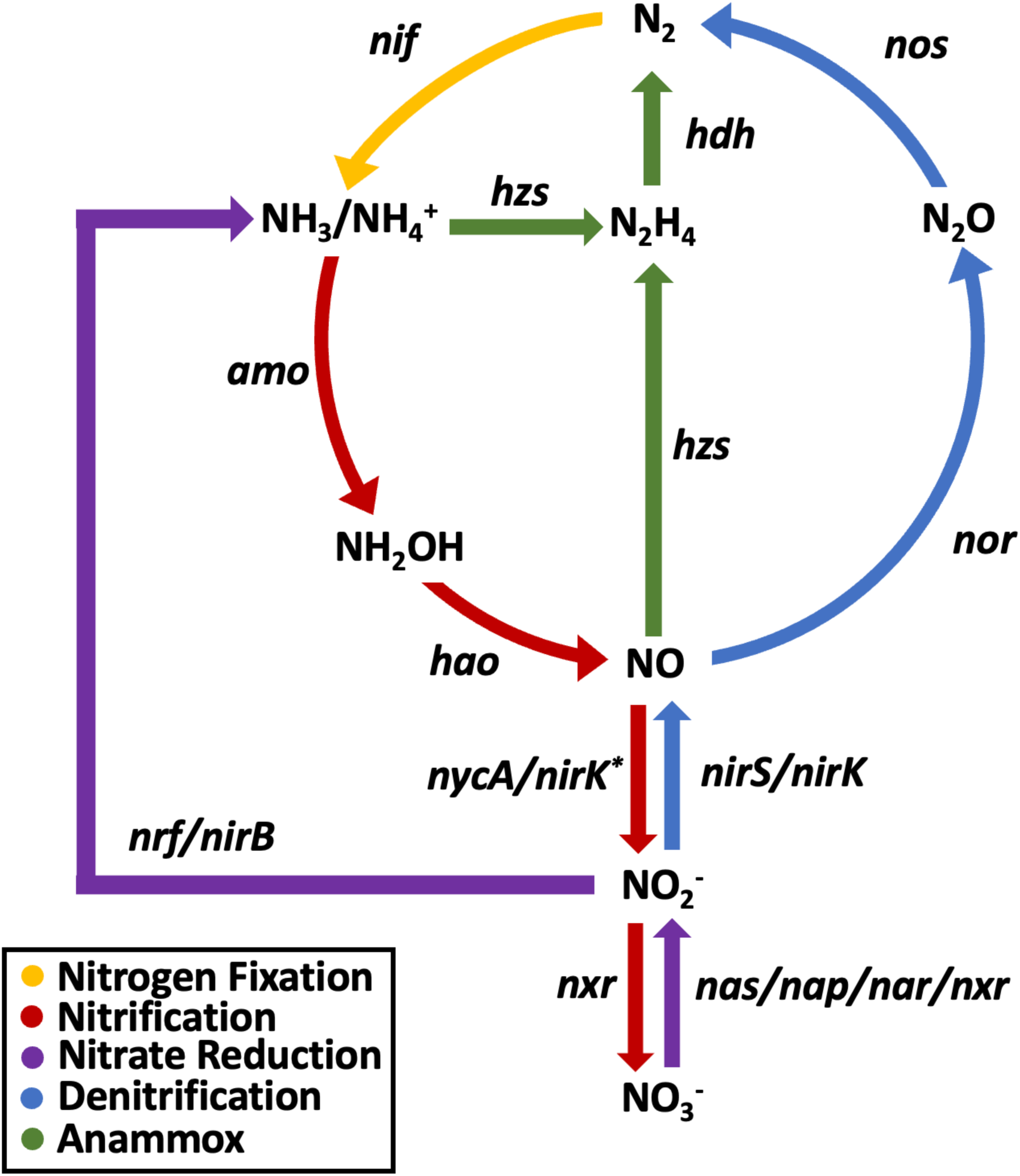
The inorganic nitrogen cycle pathway as performed by bacteria or archaea. The traditional cycle begins with the transformation of N_2_ to NH_3_ through nitrogen fixation (yellow), proceeds to NH_3_ oxidation to NO_3_^-^ through nitrification (red), and concludes with NO_3_^-^ reduction to N_2_ by denitrification (blue). Assimilatory/dissimilatory nitrate reduction (purple) and anammox (green) are specialized pathways to quickly reduce reactive nitrogen species. Gene abbreviations encode the following proteins: nitrogenase (*nif)*, ammonia monooxygenase (*amo*), hydroxylamine oxidoreductase (*hao*), nitrosocyanin (*nycA*), nitrite oxidoreductase (*nxr*), assimilatory nitrate reductase (*nas*), dissimilatory nitrate reductase (*nap/nar*), nitrite reductase (*nir/nrf*), nitric oxide reductase (*nor*), nitrous-oxide reductase (*nos*), hydrazine synthase (*hzs*), and nd hydrazine dehydrogenase (*hdh*). *Proposed genes that have not been experimentally confirmed. Adapted from (74,83).

Denitrifiers close the loop of the nitrogen cycle, reducing nitrate back to N_2_ via a series of intermediate oxidized forms including atmospheric pollutants such as NO and the greenhouse gas N_2_O (**9, 10**). Increased N_2_O emissions under nitrogen-rich conditions have been linked to increased nitrification and denitrification activity by microbes (**11, 12**). This finding raises concerns about global climate change since one ton of N_2_O has the same global warming potential as 250-300 tons of CO_2_ (**12**). An estimated 7.3 × 10^6^ tons of N yr^-1^ is released in the form of N_2_ O because of anthropogenic activity, 52% of which can be attributed to bacterial lithotrophic oxidation of excess nitrogenous fertilizer (**13**).

An improved understanding of how the microbial community contributes to the N cycle is crucial for developing fertilizers that mitigate excess emissions. Determining the composition of the microbial community that drive N cycle processes under different environmental conditions is an important first step. Despite an advance in genomic sequencing, there are still many unidentified species in both soils and freshwater, complicating models of community activity (**14**). In soils, these uncharacterized species have been identified as the drivers of nutrient cycling, making them the key to understanding the holistic community activity (**15**). For many microbiomes such as wetland and compost soils, this work is still in its infancy.

Wetlands are at the boundary of terrestrial and deeper aquatic ecosystems. Official definitions vary depending on governmental body, yet it is generally agreed that a wetland is an area that has regular flooding, wet soils for some or all of the growing season, and plants capable of sustained growth in saturated soils (**16**). Although wetlands support relatively high levels of biodiversity and play an important role in nutrient cycling, over 50% of natural wetlands in the Glaciated Interior Plains have been drained for agricultural purposes (**17**). Governmental agencies in the U.S. have recently implemented conservation and restoration programs to re-establish wetland biodiversity and improve carbon storage while creating a sink for excess NO_3_^-^ (**18, 19**). There is much study of the role of wetlands in water quality improvement via denitrification but not much on the composition of responsible microbial communities (**20–22**).

Although composting of organic material has emerged as a leading alternative to landfills, greenhouse gases (GHG) such as N_2_O are produced during decomposition (**23**). While N_2_O emissions are relatively low by mass compared to other GHGs, N_2_O has an outsized contribution to greenhouse warming potential (24). Emissions tend to be highest in the early mesophilic stage of composting, as most denitrifiers cannot survive the thermophilic stage, when trapped heat from respiration leads to selection of thermophiles (25). Ammonia emissions are also observed in well-aerated compost piles as higher temperatures promote the volatilization of NH_3_ produced during the mesophilic stage (26). To our knowledge, there has not been a thorough investigation of the complete N cycle in composts.

Shotgun metagenomic sequencing identifies novel species, such as the commamox-performing *Nitrospira*, and enables measurement of relative gene abundances in a community (27). Metagenomic sequencing is a relatively unbiased approach to characterize the entire microbial community from isolated DNA (28). After sequencing, these data can be mined for prevalent taxa and genes of interest using computational pipelines. Assembly-free techniques employ alignment-based strategies to identify the genes of interest or taxa from generated short read sequences. This marker gene approach allows us to estimate potential activity of nitrogen cycle pathways across all members of the community; however, this estimate cannot accurately model active metabolic processes since genetic material could be from dormant or deceased individuals (29, 30).

For our study, we combined existing data sets with new data gathered from wetland soils, agricultural soils, and compost to compare potential activity of N cycle pathways between diverse soil and freshwater microbiomes. Previous studies from the Slonczewski lab use assembly-free techniques to determine the relative abundance of antibiotic resistance genes as well as prevalent microbial taxa in the Kokosing River, four Knox County, OH ponds, and two lakes in the Antarctic McMurdo Dry Valleys region (31–33). We combined these datasets with new data from studies of Knox County farm soil, wetland soil, and compost. DNA collection and sequencing methods were comparable for all microbiomes.

Using NCycProfiler (34), we mined our metagenomic data for representative marker genes of the six N cycle pathways. From the complete NCyc Database (NCycDB), we selected ten highly specific markers to decrease the probability of a short read spuriously aligning to an incorrect marker sequence (**Supplemental Table 1**). We then investigated the relationship between ambient N_2_O flux in wetland soils and the denitrification gene count. Finally, we identified the most prevalent taxa from the wetland, compost, and agricultural soils with Kraken2/Bracken to determine which genera likely harbor denitrification genes (35).

## METHODS

### Site characterization for soil and compost

New metagenomic data was obtained from soil and compost collected between 2022 and 2024 in Knox County, Ohio, a scenic, agricultural community with one small city (Mount Vernon, pop. 17,000). Sources included forest soils, agricultural soils, wetland soils, and the Kenyon College Compost.

### Sampling of compost and soil

Six compost soil samples were retrieved from the Kenyon College Compost Center (40.372°N, 82.386°W) in March of 2024 to explore the microbial community and relative N cycle gene abundance. For comparison, six soil samples were collected from the nearby Kokosing Gap Trail (40.372°N, 82.386W) to explore similarities and differences between these two sampling locations. Soil was obtained from approximately 15 cm below the surface at both sites. Temperature readings for the two sites differed drastically: the Gap Trail soil averaged 6°C whereas the compost soil was 50°C. Samples were collected using a sterile chemical scoop, placed in a sterile 50-mL conical tube, and transported back to the lab for immediate DNA extraction. In the lab, approximately 200 µL of sample was added to 750 uL of DNA/RNA Shield™ and then homogenized with a bead beater for 45 minutes. The contents were then stored at -20°C overnight for DNA prep the following day.

Wetland soil samples were collected from six sites over two consecutive years (2022 and 2023) to investigate how site quality influenced relative N cycle gene abundance and microbial communities. Ballfield (40.269°N, 82.283°W) and Blackjack (40.352°N, 82.484°W) are natural wetlands with well-saturated soil and abundant vegetation. Ballfield supported skunk cabbage and cattail growth whereas Blackjack is a button-bush pool in an ash-maple forest with significant inputs of leaf litter. Kokosing (40.376°N, 82.446°W) and Batnest (40.375°N, 82.197°W) are degraded wetlands with the driest soil of the six sampled wetlands. The Kokosing wetland is part of the Kokosing River riparian buffer. Construction of a railroad and its subsequent conversion into a walking trail likely contributed to the site’s degradation. Batnest was drained and farming attempted, but due to persistent wetness it became a remnant wetland surrounded by agriculture. More recently it was incorporated as part of a public park. The Brown Family Environmental Center (BFEC, cord 40.380°N, 82.415°W) and Burtnett (40.349°N, 82.325°W) are both 25-year old restored wetlands with highly saturated soils. BFEC is a toe of slope wetland adjacent to Wolf Run Creek that receives water flow from upgradient pasture land. Burnett is a depressional wetland restored as part of the USDA Wetland Reserve Program and is surrounded by a mixture of pasture, wetland, and agricultural land.

In 2022, two 4 cm diameter by 10-cm deep cores were extracted from each site and kept on ice in Whirl-Pak bags during transport. In the lab, five to ten grams of soil were scooped from the core using sterile technique and stored at -80°C until sequencing. Cores were stored at 4°C until further analysis. Soil pH was measured with a Hannah pH/conductivity combination meter after diluting 10 g of soil in 20-mL of water for 30 minutes.

For the 2023 wetland soil sampling period, the same field protocols were followed. 5-10 g of soil were obtained using sterile forceps and placed in DNA/RNA Shield™ (Zymo Research Corp., Irvine, CA) for sample preservation.

Agricultural soil was collected from three nearby farms in 2023 according to the same field protocols. BFEC Farm (40.340°N, 82.415°W) and Burtnett Farm (40.348°N, 82.326°W) are immediately adjacent to their respective wetlands whereas McManis Farm (40.398°N, 82.415°W) is a completely distinct residential property.

### Published datasets from Ohio ponds and river and from Antarctic lakes

Published datasets for the Knox County, Ohio ponds and Kokosing river were included to increase the variety of habitat types in our analysis (31, 33). Pond and Kokosing River water was collected in Knox County, OH, a scenic agricultural community in the Ohio River watershed. Four separate ponds (Burtnett, Foundation, McManis, and Porter) were sampled over two consecutive years. All samples were filtered at 0.22 µm and processed as detailed by (31) and (33). Water pH, conductivity (𝜇S/cm), temperature (°C), and DO (mg/L) were measured as described.

Published datasets for two Antarctic lake microbiomes were also used (32). Lake Fryxell (FRY, cord 77.716°S, 162.354°E) and Lake Bonney (BON, 77.719°S, 162.372°E) are two meromictic lakes in Taylor Valley, Victoria Land, Antarctica. FRY has a maximum depth of 20 m and surface area of 7.08 km^2^ whereas BON is 40 m deep with a surface area of 4.3 km^2^. Although both lakes are covered with ∼4 m of ice year-round, biological and chemical activity differs greatly between the two sites. (36–39). Lake water was sampled in December 2014 by filtration at 0.45 µm. A sample of the Fryxell microbial lift-off mat surface was also obtained (32).

### Soil and compost DNA extraction and sequencing

DNA was extracted from the compost, forest soils, agricultural soils, and wetland soil samples according to the manufacturer’s instructions (Zymo Research Corp, Irvine, CA) as modified (33). We measured the quantity and quality of the isolated DNA using a NanoDrop One (Thermo Fisher Scientific) before shipping the extract to Admera Health (South Plainfield, NJ) for shotgun metagenomic sequencing. Libraries were prepped with Illumina® NexteraXT adapters and sequenced according to the manufacturer’s protocol (San Diego, CA) for a total of 40 million reads (20 million in each direction). We used Trimmomatic v0.39 to remove adapter regions and low quality reads from our data (40).

DNA extraction, sequencing, and short read processing for the pond, Kokosing River, and Antarctic lake water samples was performed as described previously (31–33).

### Mining Metagenomes for Nitrogen Metabolism

All metagenomes were mined for N cycle pathways using NCycDB v1.1 (34). On-target N cycle protein families and their respective annotations were identified using the KEGG ontology database (41). Amino acid sequences for these protein families were isolated from UniProt (https://www.uniprot.org/) and clustered with USEARCH v7.0 (42) at a 30% identity cutoff. Outlier sequences were then removed. To build a high-confidence, searchable database, N cycle sequences were screened against the KEGG, COG (43), eggNOG v4.5 (44), and Seed/Subsystems (45) ontology databases to identify gene families with similar sequence yet non-homologous function. Representative amino acid sequences for N cycle and off-target gene families that clustered at 95% identity or greater with CD-HIT v4.6 (46) were included in the final 95 NCycDB.

Since the NCyc pipeline only accepts a single input file, trimmed PE reads were concatenated into a single FASTQ file. We then searched our concatenated PE and Antarctica SE reads for N cycle genes using the NCycProfiler.PL script. To minimize the risk of redundancy, boost computational efficiency, and control for library size, we aligned a random subset of 5,000,000 reads from each sample to the 95 NCycDB with DIAMOND v2.0.15 (47) at an e-value cutoff of 0.001. Before alignment, DIAMOND translates the short read into its respective amino acid sequence. The total number of “hits” to each N cycle protein family was recorded.

### DEA Incubations

Denitrifying Enzyme Activity (DEA) incubations were run on the 2023 wetland and agricultural soil with ambient intact soil cores using the acetylene block technique. Each room temperature soil core, with a volume of approximately 282 mL (diameter= 6.0 cm, depth=10 cm), was weighed and then placed into a 500-mL jar sealed with a gas sampling septum, leaving 217 mL of remaining headspace. Once sealed, the headspace was evacuated and flushed with helium gas for approximately four minutes to create low oxygen conditions. The headspace was then injected with 30mL of acetylene gas. In the presence of acetylene, N_2_O is the final product of denitrification by preventing the reduction of N_2_O to N_2_. N_2_O is present in low concentrations in the atmosphere and can easily be detected by gas chromatography, making it an effective way to reliably measure denitrification rates (48). Gas samples were collected at set intervals over a 24-hour period. For each sample, 15mL of headspace was withdrawn and placed into an evacuated vial. The headspace was then replaced with 15 mL of a 1:9 acetylene:He gas mixture (49).

### Gas Chromatography

Gas samples were analyzed for N_2_O using GC-2014 Shimadzu Gas Chromatograph. N_2_O concentrations (ppm) were recorded for each sample and used to calculate ambient denitrification rates at each site. In addition, CO_2_ concentrations were recorded for all samples in order to calculate soil respiration.

### Taxonomic identification with Kraken2

The Kraken suite is a powerful, assembly-free toolkit to determine taxonomic profiles of metagenomes (35). For each of our FASTQ files, Kraken2 (v 2.1.2) first hashed the Standard RefSeq Reference Database containing archaeal, bacterial, viral, and human genome sequences (Accessed June 2023) to reduce active memory usage (50). Using a sliding window algorithm, Kraken2 then identified memory minimizing subsequences, denoted as *l*-mers, from *k*-mers of length 35 for each trimmed short read in our metagenomes. All distinct *l*-mers for a given read were then converted to a hash sequence and queried against the RefSeq hash table for key matches that correspond to the Lowest Common Ancestor (LCA). To compensate for the possibility of false key identification, the short read’s true LCA was assigned to the taxon with the greatest number of *l*-mer key matches. For each of our metagenomes, Kraken2 reported the percent abundance of taxa as a function of all sequenced reads at strain-level resolution.

The proportion of identified reads depends on read quality, reference genome completeness, and the region of genomic DNA encapsulated by the read. Therefore, a fair number of reads go unassigned, making interpretation of the Kraken2 output difficult. We employed Bracken v2.9 to calculate the relative percent abundance of genera using only Kraken2 assigned reads, improving interpretability (51). Since the LCA will often be assigned to higher taxonomy levels, Bracken implemented Bayesian probabilities to determine the likelihood that any given read should be re-classified as some lower taxon. We omitted all “human” assigned sequences because the values appeared inconsistent and increased the variability of the overall relative abundance, thus limiting the interpretation of microbial communities.

### Statistical analysis

All statistical analyses were performed with nonparametric techniques in R (v4.1.2; R Core Team 2021). Correlations between measured environmental factors and N cycle genes were determined by a Spearman correlation (52).

### Data availability

Sequence files for the Antarctica lake water, Kokosing River, and pond water are deposited under the respective NCBI BioProject numbers: PRJNA367362 through PRJNA367373, PRJNA706754, PRJNA1107813, PRJNA1283204. All custom code can be found on GitHub (https://github.com/pilatand/Honors_Biology_Project/tree/main).

## RESULTS

### Denitrification genes are more prevalent in soil than in freshwater

For the past decade, the Slonczewski lab has sequenced freshwater and soil microbiomes across Knox County, OH, and from the McMurdo Dry Valleys in Antarctica. All metagenomes were mined for nitrogen cycle marker genes using NCycProfiler (**Figure 2)** (34). The resulting heatmap of average marker gene hits is organized according to sample type (water or soil) and N Cycle pathway. Yellow-green colors indicate higher gene abundance. The soil and compost samples showed greater abundance of denitrification genes (*nirB*, *nirS*, *nirK*, *norB*, *nosZ*) than the water samples (Wilcoxon rank sum, p < 0.001). In soil samples, the Cu-containing nitrite reductase (*nirK)* was more abundant than the cd_1_-heme nitrite reductase (*nirS*, Wilcoxon rank sum, p < 0.001). Across all sites, we observed very few hits to marker genes for the nitrification and annamox pathways.

**Figure 2.**
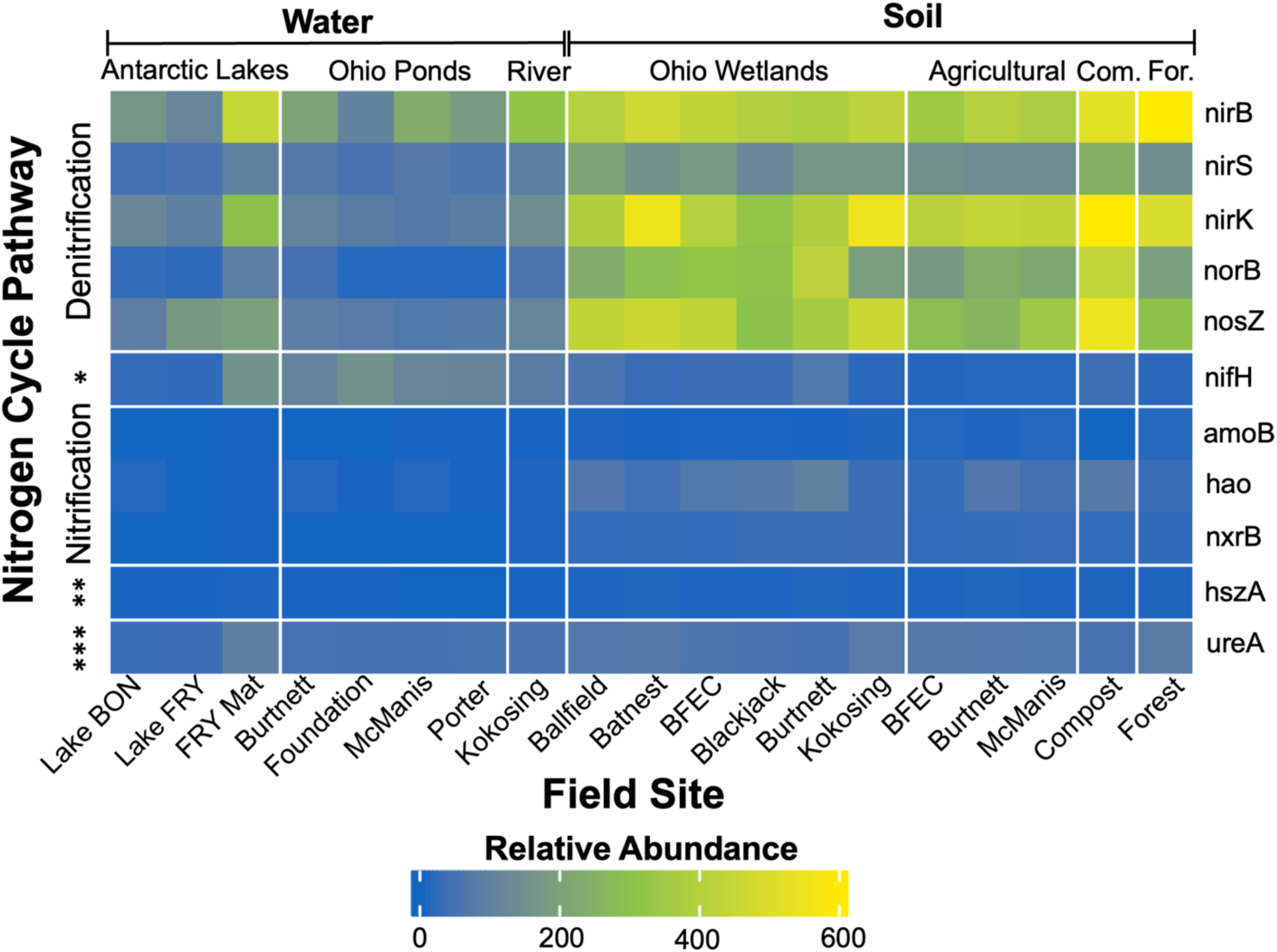
Median relative abundance of N cycle marker genes across all sampling sites. The map is clustered according to site type and metabolic pathway. Warmer colors (yellow-green) indicate that the average number of “hits” to marker genes were high while cooler colors (blue) imply the gene was less abundant. The relative abundance of assimilatory nitrate reduction and denitrification genes are significantly greater in soils (Wilcoxon rank sum, p < 0.001). N cycle gene profiles were determined with NCycProfiler. (*) Nitrogen Fixation, (**) Anammox, (***) Organic Nitrogen Decomposition.

Among the water samples, the FRY lift-off mat and Foundation Pond had the greatest number of hits on average to *norB*, the marker gene for nitric oxide reductase. These same sites also had the most hits on average for *nifH*, the marker gene for nitrogen fixation, out of all soil and water samples. The FRY lift-off mats and Foundation Pond harbor a high abundance of cyanobacteria that are capable of fixing N_2_ (32, 33). Interestingly, the FRY lift-off mat denitrification gene profile appeared more similar to the soil samples in Knox County than the Antarctic lake water. Members of the lift-off mat originate from the sediment, suggesting that benthic denitrification activity is possible despite the frigid temperatures.

Important nitrification genes including *amoB* and *nxrB* were rarely detected, which could suggest low rates of nitrification across all sites. We also detected only a few hits to hydrazine synthase (*hzsA*) in the soil and water samples. Since most samples were collected from relatively aerobic locations, it is possible that conditions were not suitable for anammox-performing bacteria, yet we find it more likely that the NCyc database struggles to identify nitrification and anammox genes from metagenomic samples. Contrary to the literature, NCycDB identified only a few hits to nitrification and anammox marker genes in groundwater (Mosley et al. 2022), implying that the pipeline underestimates the true abundance of these genes within the metagenome.

### Freshwater nitrogen cycle gene profiles associate strongly with environmental factors

The concentrations of dissolved oxygen (DO), nitrate, ammonia, and phosphate were collected for all pond and Kokosing River water samples. Using a Spearman correlation heatmap, we investigated relationships between environmental factors and the relative abundance of prominent N cycle pathways such as denitrification. Across all sampling locations, there was a significant negative correlation between DO concentration and the denitrification pathway (**Figure 3**, **Supplemental Table 2**). In the figure, yellow color indicates positive correlation whereas blue indicates negative correlation. During the 2022 pond sampling season, DO had a significant, negative correlation with *norB* (Spearman correlation, Spearman’s rho = -0.55, p < 0.001). This relationship was not observed for the Kokosing River or 2021 pond sampling periods. DO concentration also showed a significant, negative correlation with *nirB*, a gene encoding a nitrate reductase subunit, for both pond sampling periods (2021: p < 0.05, 2022: p < 0.001). This same relationship was nearly significant for the Kokosing River (p = 0.073). While the negative relationship between denitrification activity and DO concentration is well understood (54), these data provide evidence that high DO selects against bacteria capable of performing denitrification in freshwater.

**Figure 3.**
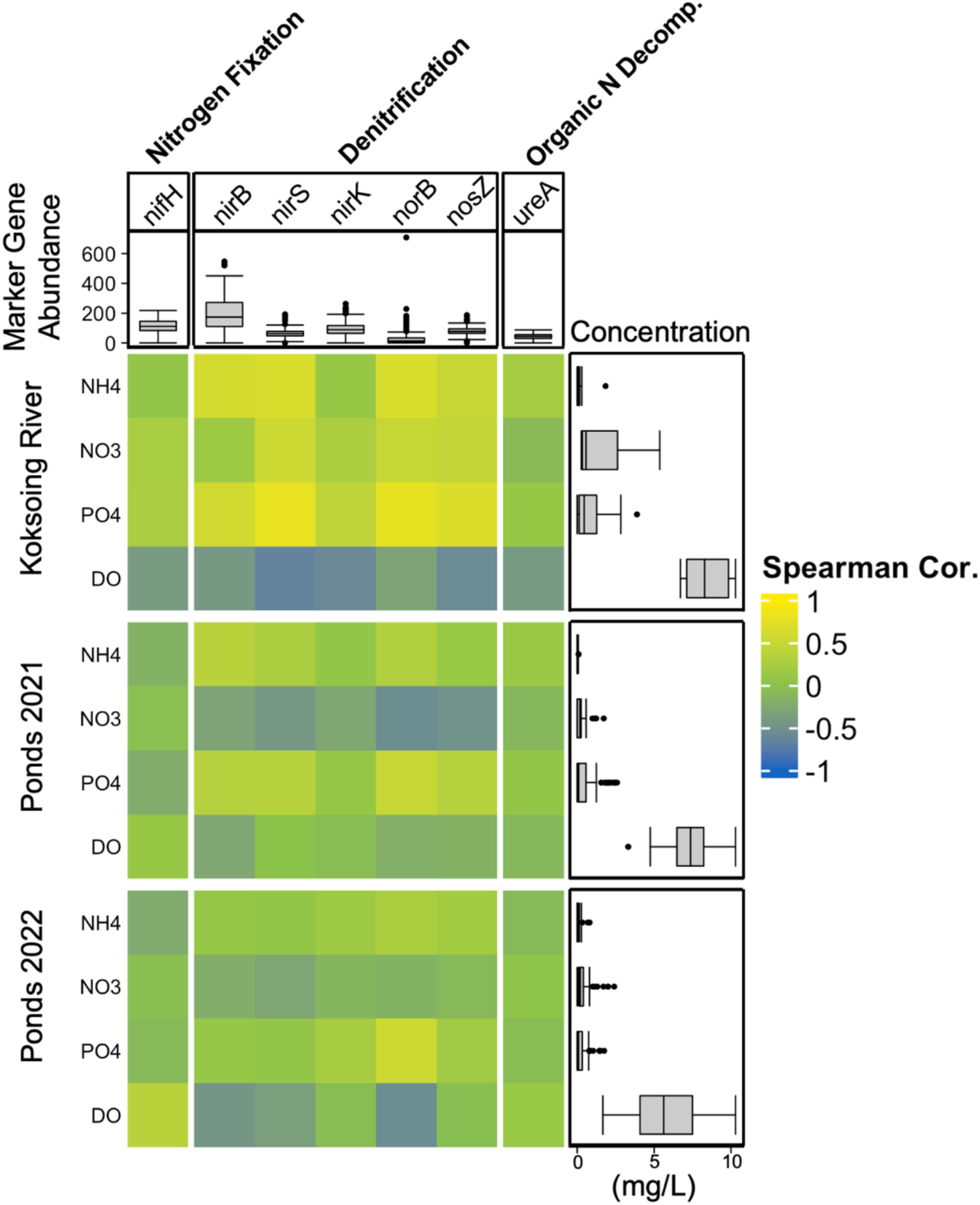
Spearman correlation heatmap of N cycle marker genes hits and pond chemistry. Phosphate shows a consistent, positive correlation with *norB* (p < 0.001). The map is clustered according to site type and metabolic pathway. Yellow color indicates positive correlation whereas blue indicates negative correlation. The distribution of N cycle marker gene hits is plotted above the map while the measured concentration (mg/L) of pond metabolites is plotted to the side. N cycle gene profiles were determined using NCycProfiler while ammonia, nitrate, and phosphate concentration was measured with Test N’ Tube Kits.

Several positive relationships were also observed between water chemistry and denitrification. All freshwater sites showed a positive association between the phosphate concentration and the denitrification pathway (**Figure 3**, **Supplemental Table 2)**. There was a strong, positive relationship between phosphate and *norB* in the Kokosing River and for both pond sampling periods (Spearman correlation, Spearman’s rho: Kokosing River = 0.78, Pond 2021 = 0.49, Pond 2022 = 0.55, All p < 0.001), whereas correlations with other denitrification marker genes varied by sampling period and site. Ammonia and nitrate were both positively associated with *nirS* and *norB* in the Kokosing River (**Figure 3**, **Supplemental Table 2**), suggesting that nitrogen input from a local wastewater treatment plant may be selecting for river denitrifiers.

### N_2_O emissions from wetland and agricultural soil correspond to relative *norB* abundance

To measure the ambient N_2_O flux in agricultural and wetland soil, we collected N_2_O gas from acetylene-inhibited cores every two hours over a 24-hour period. The average N_2_O flux was determined by fitting a regression line to the data after outlier measurements were removed. We found a clear positive association between the relative abundance of *norB* and N_2_O emissions (**Figure 4)**. The relative abundance of *norB* was significantly greater in the wetland soils than in the agricultural soils (Wilcoxon rank sum, p < 0.05). Although N_2_O flux was generally higher in the wetland soils, it was not significantly different from the agricultural soil (Wilcoxon rank sum, p = 0.231). We found no relationship between other genes in the denitrification pathway (*nirS*, *nirK*, and *nosZ*) and the N_2_O flux (**Supplemental Table 3**).

**Figure 4.**
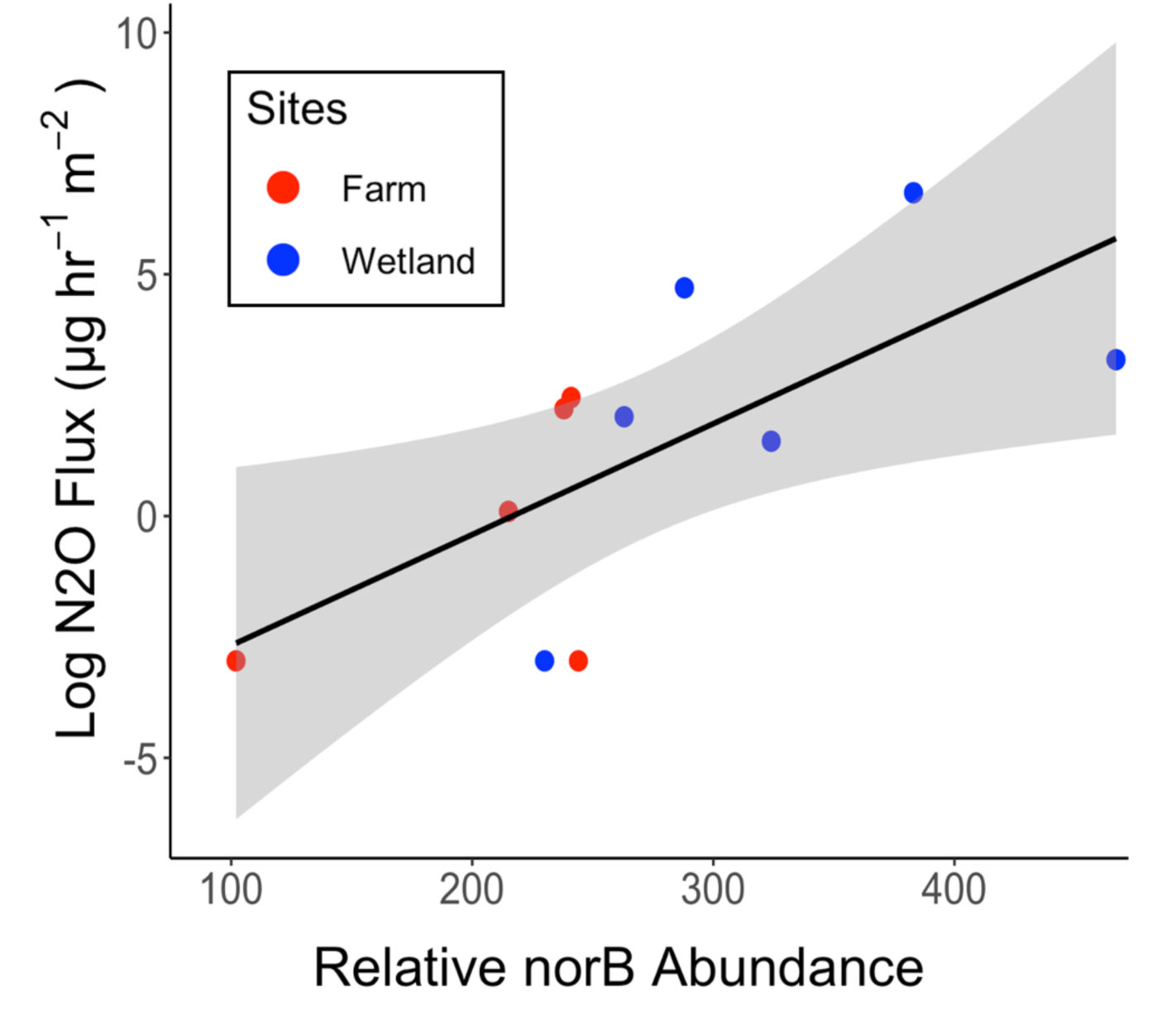
Relative abundance of *norB* predicts ambient N_2_O emissions in wetland and agricultural soils. (p = 0.023, R^2^ = 0.396). The relative abundance of *norB* was determined from sequenced metagenomes with NCycProfiler. The average rate of N_2_O flux was determined over 8 hours from acetylene-inhibited soil samples.

### Kraken2/Bracken taxa profiles from soils reveal high abundance of denitrifier genera

Our marker gene analysis indicates that the Kenyon Compost and soil samples harbor many microbes capable of performing denitrification, and their abundance is much higher than in corresponding water samples. To identify these genera, we mined our metagenomes with the Kraken2/Bracken pipeline (35).

We compared the community composition of wetland soils with agricultural soils collected only a few meters away. Across all sites, there was no noticeable difference in the relative abundance of Alphaproteobacteria, Actinomycetota (or Actinobacteria), and Betaproteobacteria species (**Supplemental Figure 1**). *Bradyrhizobium* were the most dominant of the Alphaproteobacteria, representing over 5% of the identified microbial community in most of the soil samples. Although *Bradyrhizobium* species are primarily known for nitrogen fixation on legumes, the denitrification pathway is energetically favored in several members such as *Bradyrhizobium denitrificans* and *Bradyrhizobium japonicum* when carbon sources are scarce (55–58). In all samples labeled “Ohio Wetland Soils” (**Figure 2**), *Bradyrhizobium* was strongly associated with *norB*, suggesting that they could be the primary drivers of denitrification in these anoxic soils (Spearman correlation, 𝜌 = 0.62, p < 0.001).

### Compost exhibits high abundance of denitrifying thermophiles

Thermophilic microorganisms are common in composts following the oxidation of simple sugars in the mesophilic stage (23). Compared to the soil collected from the nearby Kokosing Trail, the compost exhibited a high prevalence of Thermomicrobia, a class of bacteria containing two thermophilic genera (**Figure 5, Supplemental Figure 2**). *Sphaerobacter* were the most common Thermomicrobia MAG assembly of *Sphaerobacter thermophilus*, the only known member of the *Sphaerobacter*, has revealed that the species harbors several denitrification genes such as *nirK* and *nosZ* (59, 60). We also identified several genera of methanogenic archaea that were unique to the compost samples. Whereas members of the *Methanosarcina* such as *Methanosarcina thermophila* and *Methanosarcina bakeri* have been previously characterized in composts, this is the first study to find *Methanocella* and *Methanotrhix* species (61). Thermophilic members of these genera have been found in sludge, natural wetlands, and rice fields, suggesting that composts provide suitable conditions for their growth (62, 63).

**Figure 5.**
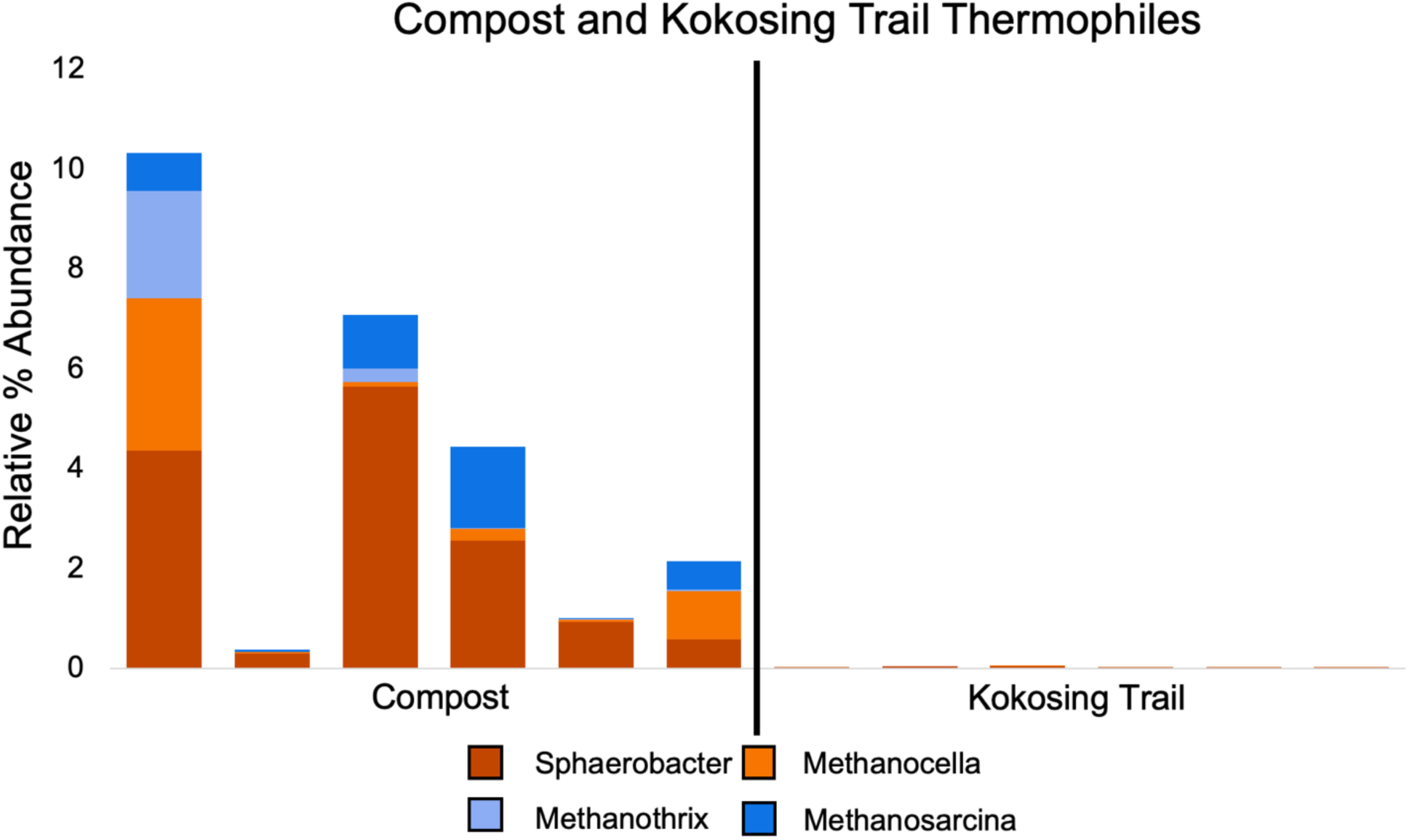
Thermophilic bacteria and methanogenic archaea are more prevalent in compost than nearby Kokosing Trail soil. *Sphaerobacter* species from the bacterial class Thermomicrobia and several archaeal methanogens thrive in the thermophilic compost. The relative percent abundance at the genus level was determined by the Kraken2/Bracken pipeline.

## DISCUSSION

As global temperatures continue to rise, it has become increasingly necessary to characterize how ecosystems contribute to natural greenhouse gas emissions. The nitrogen cycle is driven by microbial activity; therefore, improved understanding of environmental factors that select for N_2_O emitting bacteria is required before mitigation strategies can be designed. To determine potential nitrogen cycle processes in six distinct microbiomes, we used an assembly-free, metagenomics-based approach. We then analyzed how environmental factors select for nitrogen cycle genes in the Kokosing River and four Knox County ponds. Since the gene profile alone does not provide information about which taxa are responsible for the different nitrogen cycle pathways, we also identified the most prevalent genera in these microbiomes with the Kraken2/Bracken pipeline. We identified several genera such as *Bradyrhizobium* that are likely responsible for denitrification activity in these microbiomes.

### NCyc identifies denitrification genes but struggles with other pathways

Using archival metagenomic data from the Slonczewski lab, we identified potential nitrogen cycle processes in six distinct microbiomes with the NCyc pipeline. Nitrogen gene profiles were found to be clustered according to sample type — soil or water — regardless of the sampling location. According to NCycProfiler, several denitrification genes such as *nirK*, *norB*, and *nosZ* were more prevalent in the soil than in the water samples (**Figure 2**). In our analysis, lift-off mats from Lake Fryxell (FRY) were grouped with water samples since the biofilms assembled under the ice; however, the nitrogen gene profile of the oxygenic, top layer appeared more like the Ohio soil samples. This is consistent with a previous study into microbial mats of the Miers Lake in the McMurdo Dry Valley that found N_2_ and CO_2_ fixation activity can reach rates similar to what is observed in soils across the continental United States despite the extreme environment (64). Although denitrification genes have been identified in the benthic region of the FRY mats, our study provides some of the first evidence that the Antarctic mat surface may be contributing to N_2_O emissions (65).

Across all sites, genes corresponding to the bacterial nitrification and anammox pathways were not commonly found (**Figure 2**). Considering that nitrification is a ubiquitous process in soils, we anticipated that these samples would have a sizeable number of hits to ammonia monooxygenase (*amo)* and nitric oxidoreductase (*nxrB*) (66). While it was not the focus of our study, NCyc also reported few hits for *amo* in ammonia-oxidizing archaea, the primary drivers of nitrification in soils (67). It was recently discovered that AMO produces N_2_O (68, 69). Thus, our approach is clearly missing a critical contributor to overall N_2_O emissions from each microbiome.

We were also surprised that NCyc only recorded a few hits to *hzsA*, the marker gene for hydrazine synthase, since microbes capable of performing anammox are commonly found in waste water treatment plants (WWTPs), soils, and freshwater (70). Whereas the 95% identity NCycDB contained hundreds of reference sequences for denitrification marker genes, nitrification and anammox genes often had fewer than 50, suggesting the tool is limited by the number of reference sequences. This is likely because NCycDB is trained on published amino acid sequences with high similarity, meaning that the pipeline likely ignores reads generated from poorly annotated species (35). Further, proteins with homologous function but non-homologous structure are also likely filtered out early in database construction. We thus conclude that NCycDB v1.1 is biased toward the denitrification pathway.

Freshwater and soil are two major areas where current metagenomic techniques fail to capture true microbial diversity and metabolic activity; therefore, tools like NCyc will likely become more accurate as our understanding of the so-called microbial dark matter — species that are currently unculturable — improves (14).

### Water chemistry corresponds to potential denitrification activity

For the freshwater sites, DO was negatively correlated with denitrification genes (**Figure 3**). The Kokosing River and pond water were collected from the epilimnion where DO concentration varied substantially (2-11 mg/L). Although denitrification in freshwater is commonly performed in the sediment where oxygen content is minimal, some activity can be measured in the aerobic water column, going against the traditional model of denitrification (71). When oxygen is present, it is assumed that aerobic respiration will be preferred, limiting the necessary electrons for denitrification; however, a mechanism under slightly aerobic conditions — perhaps by bacteria embedded in detritus or biofilm — has been proposed by (72). These authors assembled an artificial, oxygenated freshwater system with sufficient nitrate for denitrification. They fixed an iron rod into their sediment, hypothesizing that freshwater bacteria would mediate electron transfer. After a few days, denitrification activity could be measured, driven by a unique biofilm that accumulated on the rod containing *Dechloromonas*, *Simplicispira*, and *Hydrogenophaga* species. It is important to note that this mechanism has not been confirmed in the field. Considering that *Hydrogenophaga* and *Dechloromonas* species were commonly observed in our freshwater samples and DO was a significant predictor of the community composition (31, 33), it is possible that denitrification is being performed in anaerobic microenvironments within the slightly aerobic water column. When DO concentration is high, denitrification is likely completely inhibited.

Interestingly, we also observed a consistent, positive relationship between denitrification genes and the PO_4_ concentration in the freshwater samples (**Figure 3**). High phosphate input from fertilizers is directly linked to harmful algal blooms (HAB) in lakes even when forms of reactive N are absent. Identifying mechanisms that can sufficiently remove phosphate from the environment has thus become a top priority for those seeking to reduce the likelihood of HAB (73). Since natural sources of phosphorus are anticipated to be depleted within the next 100 years, focus has shifted toward discovering microbial pathways that can capture the metabolite before it is permanently lost to the ocean (74, 75). Denitrifying phosphorus removal has become an attractive option as nitrate and phosphorus uptake are coupled under anoxic conditions (76). These same conditions also happen to favor HAB, implying that denitrifying phosphorous removal may a primary driver of increased N_2_O emissions in lakes during a bloom. Although conditions were not drastic enough to cause a bloom, it is probable that some denitrifying phosphorus removal was being performed. Regardless, the introduction of harmful metabolites and pathogenic genera from the WWTP effluent such as *Acinetobacter* and *Pseudomonas* indicate that wastewater increased the risk of HAB and severe infection in late 2019 and early 2020.

### Ambient N_2_O flux as a function of gene content offers promising area for future study

Agricultural activity and natural soils collectively emit 9.4 Tg N_2_O yr^-1^, making them a key target for emissions reduction (13). Historically, wetlands have been drained for supposedly more productive purposes, including land development and farming (77) which increase N_2_O emissions. A recent meta-analysis of global wetland restoration programs on agricultural lands provided strong evidence that rehabilitation results in substantial reduction of measurable N_2_O (78).

Since microbes drive denitrification in wetland soils, we hypothesized that the metagenome might serve as a reliable predictor of N_2_O emissions. We found that ambient N_2_O flux correlated strongly with *norB*, the marker gene for nitric oxide reductase, yet there was no obvious relationship between N_2_O flux and site condition (**Figure 4**). A study of 72 wetlands across the globe identified ammonia-oxidizing archaea as the primary drivers of N_2_O flux (79). The researchers only employed metagenomes to determine the most prevalent bacteria and archaea in their samples. They used PCR amplification to identify the relative abundance of *nirS*, *nirK*, and *nosZ* sequences in their samples, limiting their possible conclusions. Nonetheless, our findings concur with their conclusion that despite our detected *norB* relationship there is no significant relationship between the other three denitrification genes and N_2_O (**Supplemental Table 2**).

Another study considered N_2_O flux from soils collected from four experimental farms in France. Again, the team used PCR amplification of the most common denitrification marker genes — *nirS*, *nirK*, and *nosZ* — to understand what conditions in agricultural soils are responsible for N_2_O emissions (80). They found that soil properties such as pH were the best predictors of *in situ* emissions whereas the selected marker genes had no significant relationship. Given the modularity of denitrification, we believe that a comprehensive analysis of all genes in the pathway is necessary to accurately predict ambient N_2_O flux. Future study of wetland and agricultural soils should consider *norB* as a predictor of flux since it plays such an important role in the actual production of N_2_O. It would also be of interest to investigate if the relative abundance of *nosZ* can predict N_2_ flux from soil.

### Pipeline determination: a critical choice for metagenomics

Although it is widely accepted that assembly of contiguous sequences can improve the accuracy of gene and taxa identification, assembly may introduce many errors if the workflow is not properly configured, resulting in a plethora of artificial MAGs (81). To minimize the likelihood of introducing error, we used two alignment-based, assembly-free algorithms to mine our metagenomes for genes and taxa of interest: Kraken2/Bracken, a workflow for identifying microbial taxa (35) and NCycDB, a tool that matches short reads to a representative database of published N cycle amino acid sequences (34).

One major drawback of using Kraken2/Bracken is its limited taxa resolution. Although one can theoretically reassign abundance estimates down to the species level, this method obscures clear, consistent trends. Our current approach allows us to make high confidence hypotheses about which species harbor certain N cycle genes, yet we cannot make a definitive conclusion. One solution is to bin the short reads according to species with a tool like BBsplit, a component of the Joint Genome Institute’s BBTools toolkit (82). If our proposed hypotheses are correct, then we would expect a subset of reads assigned to a particular species to account for a large proportion of hits to some marker gene. This would allow us to confirm if *Bradyrhizobium* species are truly responsible for potential denitrification activity in their respective microbiomes.

Although metagenomics can tell us a lot about how environmental factors select for or against prevalent genera, we cannot draw conclusions about how these same factors influence actual metabolic activity; therefore, future study of the N cycle in these microbiomes should incorporate robust sampling of the metatranscriptome and metaproteome.

## ACKNOWLEDGEMENTS

This Project was funded by National Science Foundation grant MCB-1923077 and by the Kenyon Philip and Sheila Jordan Fund. We thank the authors of Murphy et al. 2021, Tallada et al. 2023, Vaccaro et al. 2025, and the Kenyon College Spring 2024 Microbiology class for help with data collection.

## Notes

### Competing Interest Statement

The authors have declared no competing interest.

