## Supplemental Figures and Tables for "Comparative Analysis of Denitrification and Other Nitrogen Cycle Genes in Diverse Environmental Microbiomes"

### Supplemental Files for Comparative Analysis of Denitrification and Other Nitrogen Cycle Genes in Diverse Environmental Microbiomes

By

Andrew M. Pilat,[a] Rachael E. Tomasko,[a] Audrey Gibson,[a] Daniel Barich,[a] Zachary Slimak,[a] M. Siobhan Fennessy,[a] and Joan L. Slonczewski[a]*

[a]Department of Biology, Kenyon College, Gambier, OH

*Corresponding author: Joan L. Slonczewski, Department of Biology, Kenyon College, Gambier, OH

### Table of Contents


| | |
|---|---|
| Supplemental Table 1 | 3 |
| Supplemental Table 2 | 5 |
| Supplemental Table 3 | 6 |
| Supplemental Figure 1 | 7 |
| Supplemental Figure 2 | 9 |

**Supplemental Table 1.** Nitrogen cycle reactions as performed by bacteria and archaea. Nitrate reduction refers to the assimilatory and dissimilatory pathways. When applicable, the marker gene is included in parentheses. Adapted from Kuypers et al. (2018) and Mobley et al. (1995).

| Pathways | Bacteria | Enzyme (Marker Gene) | Reaction |
|---|---|---|---|
| *Nitrogen Fixation* | Nitrogen Fixers | Nitrogenase (*nifH*) | $N_2 + 8e^- + 8H^+ + 16\,ATP \rightarrow 2NH_3 + H_2 + 16ADP + 16P_i$ |
| *Nitrification* | Ammonia Oxidizing | Ammonia Monooxygenase (*amoB*) | $NH_4^+ + O_2 + 2e^- + H^+ \rightarrow NH_2OH + H_2O$ |
| | | Hydroxylamine Oxidoreductase (NA) | $NH_2OH \rightarrow NO + 3e^- + 3H^+$ |
| | | Nitric Oxide Oxidoreductase (NA) | $NO + H_2O \rightarrow NO_2^- + e^- + 2H^+$ |
| | Nitrite Oxidizing | Nitrite Oxidoreductase (*nxrB*) | $NO_2^- + H_2O \rightarrow NO_3^- + 2e^- + 2H^+$ |
| *Nitrate Reduction* | Various | Nitrate Reductase (NA) | $NO_3^- + 2e^- + 2H^+ \rightarrow NO_2^- + H_2O$ |
| | | Nitrite Reductase (NA) | $NO_2^- + 6e^- + 8H^+ \rightarrow NH_4^+ + 2H_2O$ |
| *Denitrification* | Denitrifiers | Nitrate Reductase (*nirB*) | $NO_3^- + 2e^- + 2H^+ \rightarrow NO_2^- + H_2O$ |
| | | Nitrite Reductase (*nirS* and *nirK*) | $NO_2^- + e^- + 2H^+ \rightarrow NO + H_2O$ |

**Supplemental Table 1.** Cont.

| Pathways | Bacteria | Enzyme (Marker Gene) | Reaction |
|---|---|---|---|
| *Denitrification (Cont.)* | | Nitric Oxide Reductase (*norB*) | $2NO + 2e^- + 2H^+ \rightarrow N_2O + H_2O$ |
| | | Nitrous-oxide Reductase (*nosZ*) | $N_2O + 2e^- + 2H^+ \rightarrow N_2 + H_2O$ |
| *Anammox* | Anammox | Nitrite Reductase (NA) | $NO_2^- + e^- + 2H^+ \rightarrow NO + H_2O$ |
| | | Hydrazine Synthase (*hzsA*) | $NO + NH_4^+ + 3e^- + 2H^+ \rightarrow N_2H_4 + H_2O$ |
| | | Hydrazine Dehydrogenase (NA) | $N_2H_4 \rightarrow N_2 + 4e^- + 4H^+$ |
| *Organic N Decomposition* | Ureolytic | Urease (*ureA*) | $CH_4N_2O + H_2O \rightarrow 2NH_2 + H_2CO_3$ |

**Supplemental Table 2.** P-values for Spearman correlation heatmap of N cycle genes and environmental factors (* = p <0.05, ** = p < 0.01, *** = p <0.001).

| Kokosing | *nifH* | *nirB* | *nirS* | *nirK* | *norB* | *nosZ* | *ureA* |
|---|---|---|---|---|---|---|---|
| NH4 | 0.819 | 0.007** | 0.004** | 0.678 | 0.004** | 0.038* | 0.436 |
| NO3 | 0.347 | 0.527 | 0.0181* | 0.299 | 0.046* | 0.051 | 0.724 |
| PO4 | 0.307 | 0.010* | <0.001*** | 0.083 | <0.001*** | 0.002** | 0.662 |
| DO | 0.904 | 0.073 | 0.003** | 0.009** | 0.195 | 0.017* | 0.084 |

| Pond 2021 | *nifH* | *nirB* | *nirS* | *nirK* | *norB* | *nosZ* | *ureA* |
|---|---|---|---|---|---|---|---|
| NH4 | 0.162 | <0.001*** | 0.027* | 0.563 | 0.008** | 0.318 | 0.288 |
| NO3 | 0.735 | 0.006** | <0.001*** | 0.014* | <0.001*** | <0.001*** | 0.312 |
| PO4 | 0.048* | 0.002* | 0.002** | 0.525 | <0.001*** | 0.004** | 0.577 |
| DO | 0.356 | 0.018* | 0.993 | 0.614 | 0.096 | 0.130 | 0.306 |

| Pond 2022 | *nifH* | *nirB* | *nirS* | *nirK* | *norB* | *nosZ* | *ureA* |
|---|---|---|---|---|---|---|---|
| NH4 | 0.096 | 0.570 | 0.578 | 0.330 | 0.068 | 0.282 | 0.400 |
| NO3 | 0.938 | 0.083 | 0.021* | 0.204 | 0.115 | 0.336 | 0.839 |
| PO4 | 0.173 | 0.078 | 0.168 | 0.028* | <0.001*** | 0.033* | 0.966 |
| DO | <0.001*** | <0.001*** | 0.002** | 0.627 | <0.001*** | 0.677 | 0.313 |

**Supplemental Table 3.** Spearman correlation p-values for denitrification genes and flux data. (* = p <0.05, ** = p < 0.01, *** = p <0.001)

| | *nirS* | *nirK* | *norB* | *nosZ* |
|---|---|---|---|---|
| Log Flux (µg $hr^{-1}$ $m^{-2}$) | 0.336 | 0.570 | 0.017* | 0.175 |

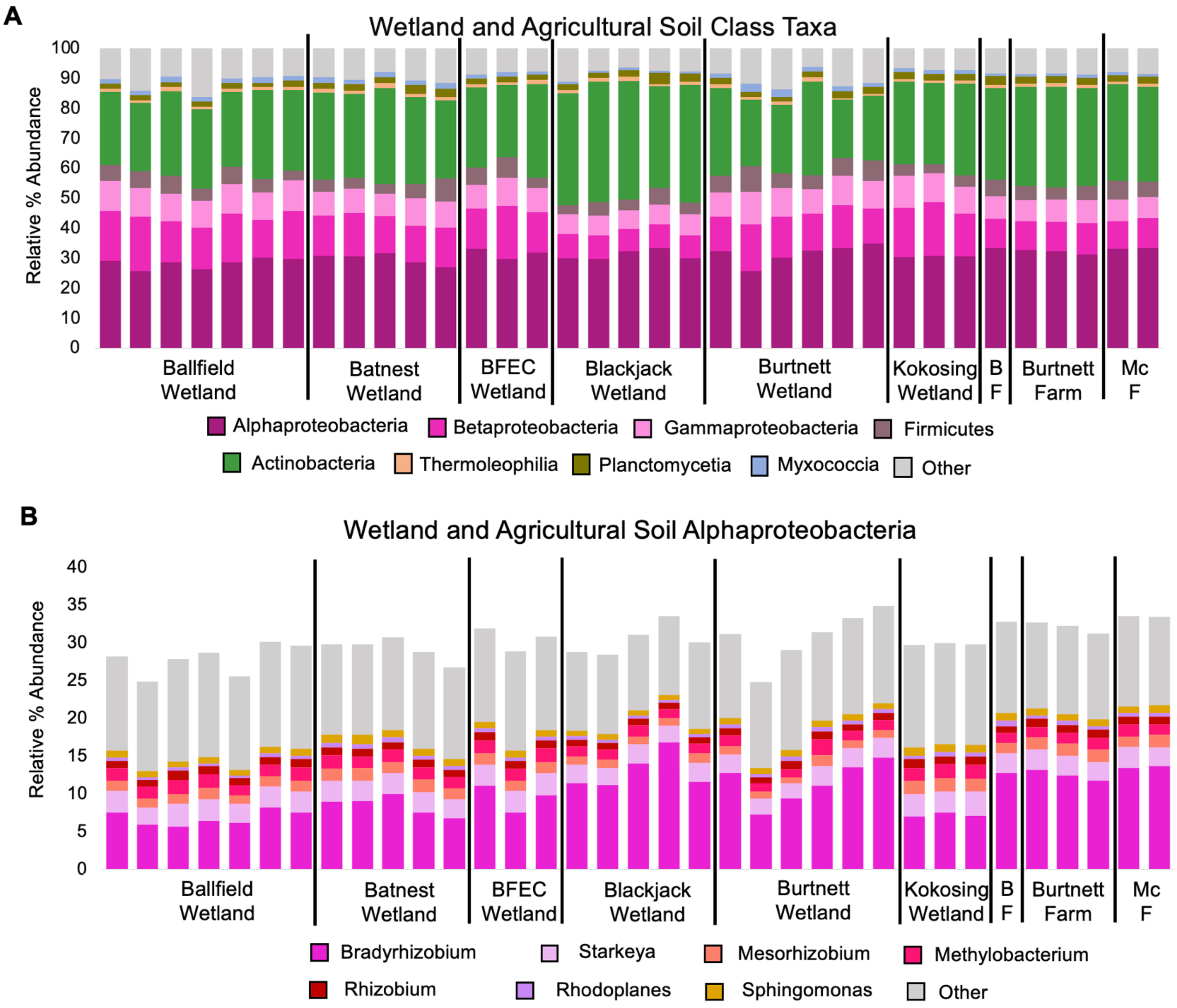


**Supplemental Figure 1. Blackjack wetland shows high prevalence of *Mycobacterium* species.** Most prevalent taxa at the (**A**) phylum and class level were the (**B**) Alphaproteobacteria determined by the Kraken2/Bracken pipeline. BF = BFEC Farm and MC F = McManis Farm.

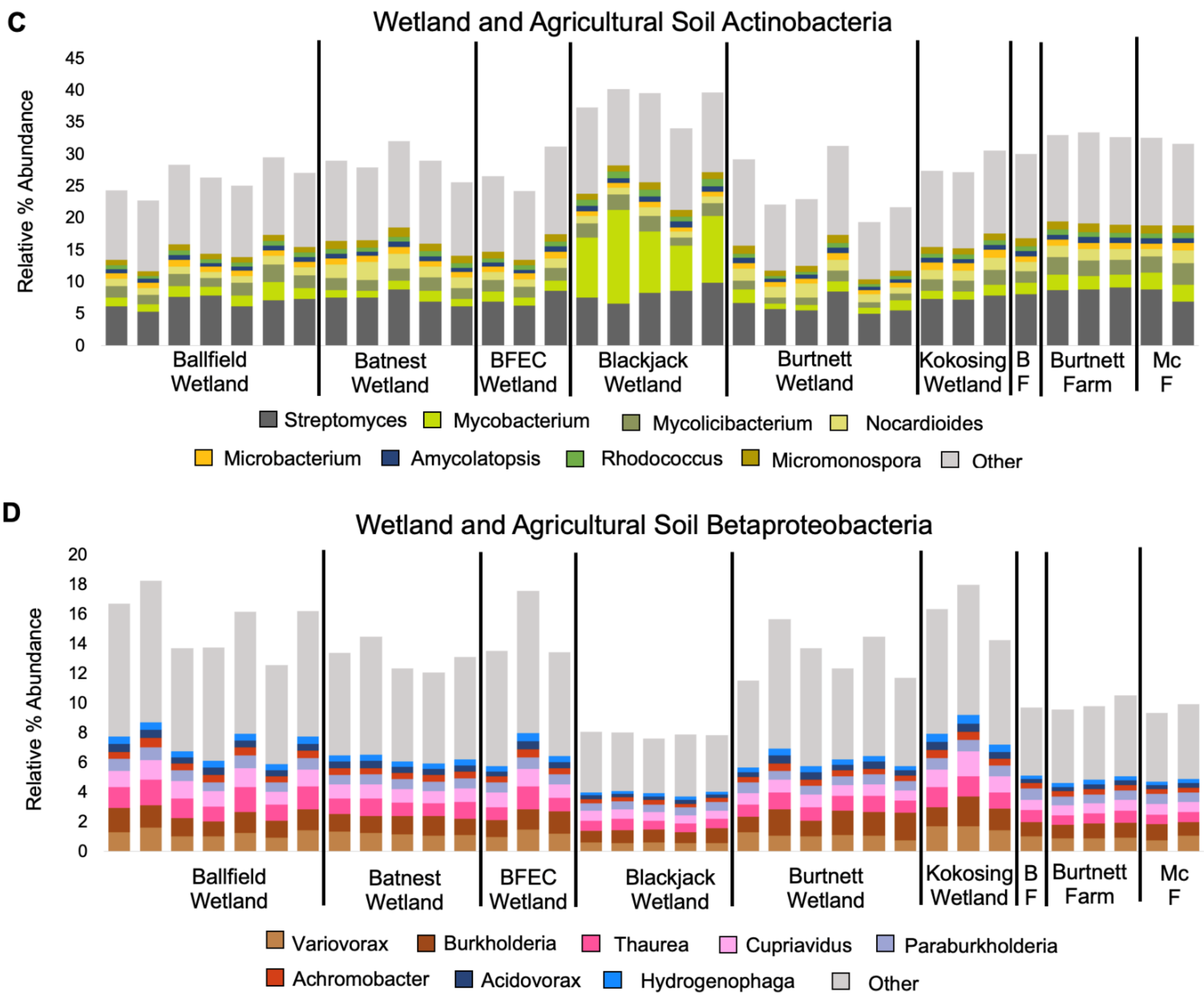


**Supplemental Figure 1 Cont. Blackjack wetland shows high prevalence of *Mycobacterium* species.** Most prevalent taxa at the class level were (**C**) Actinobacteria, and (**D**) Betaproteobacteria as determined by the Kraken2/Bracken pipeline. BF = BFEC Farm and MC F = McManis Farm.

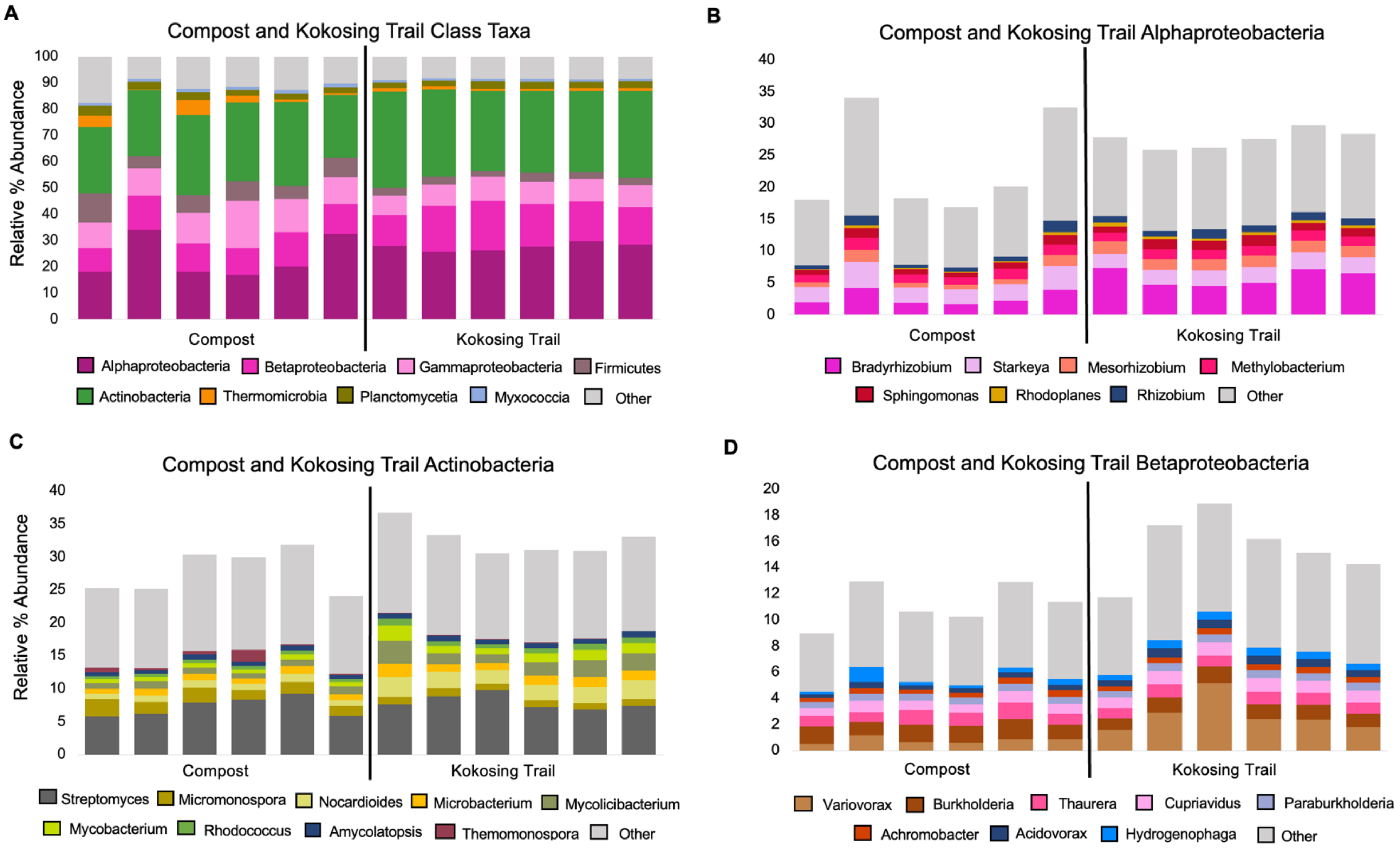


**Supplemental Figure 2. Compost samples contain thermophiles not found in nearby soil samples.** The relevant percent abundance of prevalent taxa was identified from metagenomes with the Kraken2/Bracken pipeline. Most prevalent taxa at the (**A**) phylum and class level were the (**B**) Alphaproteobacteria, (**C**) Actinobacteria, and (**D**) Betaproteobacteria.
